# Development of a Norway Rat Hepacivirus Reporter for High-Throughput Quantification of Neutralising Antibodies

**DOI:** 10.1101/2025.11.13.688247

**Authors:** Matthew J. Kennedy, Raphael Wolfisberg, Caroline E. Thorselius, Laura Collignon, Emma A. Lundsgaard, Louise Nielsen, Kenn Holmbeck, Jens Bukh, Troels K. H. Scheel

## Abstract

Hepatitis C virus (HCV) remains a major global health challenge, with vaccine research hindered by lack of immunocompetent animal models. Infection with the HCV-related Norway rat hepacivirus 1 (NrHV) in rats or mice shares HCV defining characteristics, including chronicity, delayed immune responses, and liver disease, thereby providing immunocompetent small animal models for immune responses and pathology. Infectious NrHV cell culture systems allow studies of host factors and neutralising antibodies, however, quantification relies on laborious manual counting of infected cells. To permit high-throughput quantification, we engineered culture adapted NrHV to encode green fluorescent protein (GFP), firefly luciferase (FLuc), or the 11 amino acid HiBiT tag at sites corresponding to those permissive for HCV. While GFP and FLuc constructs were not viable, a reporter with a HiBiT insertion in NS5A domain III was replication competent and produced infectious virus in rat hepatoma cells. Two mutations in NS5A adapted this virus to levels comparable to wildtype, enabling robust luciferase readouts and stability. The NrHV-HiBiT-NS5A reporter was viable in rats and permitted quantification of luciferase from liver homogenates. Luciferase-based quantification of viral infection after antibody neutralisation correlated well with manual quantification. High-throughput luciferase quantification was also used to confirm dependency on the known host factors SR-BI and miR-122, and to assess sensitivity to antivirals. Finally, sensitivity to a specific inhibitor suggested that cyclophilin A, similarly to HCV, may also be a host factor for NrHV. This novel reporter system could accelerate characterisation of the NrHV model as well as ongoing vaccine studies for HCV.

**Importance:** Direct acting antiviral therapy provides an effective cure for hepatitis C virus (HCV) infection, but the infection burden remains high due to cost, availability of therapy, and low diagnosis rates. A vaccine therefore is critical for global control of transmission, however development is constrained by a lack of immunocompetent animal models. Norway rat hepacivirus 1 (NrHV) infection of rats and mice serves as an effective small animal model for HCV, permitting evaluation of vaccine platforms and candidates. Neutralising antibody responses are considered crucial in protection and can be quantified using the NrHV cell culture system, however, only through laborious low-throughput methods. We here present an NrHV reporter virus allowing high-throughput quantification of antibody neutralisation, virus-host interactions, and antiviral efficacy via luciferase readout. This reporter will facilitate characterisation of NrHV and support ongoing vaccine research with the potential for global control of HCV.

## Introduction

Hepatitis C virus (HCV) chronically infects ∼50 million people worldwide leading to increased risk of liver cirrhosis and hepatocellular carcinoma (1). While direct acting antiviral (DAA) therapy achieves cure rates of >90%, this remains insufficient for global control of the virus due to low diagnosis rates and the high cost and low accessibility of DAA therapy in low- and middle-income countries with the highest prevalence of HCV infection (2, 3). Accordingly, a vaccine is required to achieve global control of HCV (4).

Vaccine research for HCV, which naturally only infects humans and experimentally also chimpanzees, suffers from lack of immunocompetent animal models for challenge studies. While small animal models, such as entry factor transgenic or human liver-chimeric mice, are available, they are challenged by lack of robust infection or blunted immune responses respectively, making them unsuited for use in studies of vaccines and immunopathology (5). For years, the closest known HCV relative was GB virus B (GBV-B), a virus infecting New World monkeys (6). Deep sequencing efforts have more recently resulted in the expansion of our knowledge of the hepacivirus genus, revealing HCV-related viruses in a variety of species including rodents, bats, monkeys, horses, and cattle (7, 8). Among these, Norway rat hepacivirus 1 (NrHV; rodent hepacivirus of *Rattus Norvecigus*, RHV-rn1), is of particular interest as an HCV model given its tropism for small rodent laboratory animals.

Similarly to HCV, NrHV is a positive-stranded RNA virus of ∼9.6 kb, with a single open reading frame (ORF) flanked by 5’ and 3’ untranslated regions (UTRs). Translation is driven from an internal ribosomal entry site (IRES) in the 5’ UTR and leads to expression of a single polyprotein that is predicted to be processed into three structural (Core, E1, and E2) and seven non-structural (p7, NS2, NS3, NS4A, NS4B, NS5A, and NS5B) proteins. NrHV furthermore relies on the liver specific microRNA, miR-122, for replication (9, 10). NrHV entry into hepatocytes depends on rat orthologs of the HCV host factors scavenger receptor class B type 1 (SR-BI), CD81, and occludin (OCLN), as well as claudin-3 (CLDN3), although there may be redundancy in the use of some of these factors (11, 12). NrHV leads to chronic, high titre, hepatotropic infection in rats, inducing a delayed development of specific T-cell and neutralising antibody (nAb) responses, and with the capacity to induce liver damage (11, 13, 14). With hallmarks of infection comparable to those of HCV, NrHV is well suited as an immunocompetent model for the study of hepaciviral infection, pathology, and immune responses. NrHV also infects mice, in which infection is acutely resolved within 2-3 weeks. Infection can be extended to 5 weeks through mouse-adaptive mutations or even to chronicity via depletion of CD4^+^ T-cells preceding infection (10, 15). The development of NrHV replicon and infectious cell culture systems in rat hepatocytes permitted the study of the full viral life cycle *in vitro*, providing means to study virus-host interactions and to assess and quantify nAbs generated from infected animals (9, 11). The variety of *in vivo* and *in vitro* systems available for NrHV thus make it a compelling model for essential research into pathology, immune responses, and putative vaccine candidates and combinations thereof in the ongoing quest for an HCV vaccine (7).

The current culture systems allow studies of host factors, antiviral compounds, and NrHV targeting nAbs (11, 15–17), however, they rely on laborious manual quantification of focus forming units (FFUs) following antigen staining. An NrHV reporter virus would permit high-throughput analysis of nAbs and host factors with reduced inter-operator variation. We here present the development of a viable and stable NrHV reporter virus by the insertion of the HiBiT tag (18) and accompanying adaptive mutations into NS5A domain III. Luciferase signal from the HiBiT detection system led to quantification of viral infection after antibody neutralisation, host factor inhibition, or antiviral compounds comparable to manual FFU counts, but in a high-throughput manner. This provides a much-needed tool to expedite vaccine research using the NrHV model. In addition, this reporter could aid further characterisation of NrHV, increasing our knowledge of hepacivirus evolution and the utility of NrHV as a model.

## Results

### Low permissiveness for insertion of reporter genes into the NrHV ORF

To generate an NrHV reporter virus, we first inserted a UpA/CpG-optimised firefly luciferase (FLuc) gene (19) into the ORF of RHVcc-1, a cell culture adapted clone of the RHV-rn1 strain (11). Insertion sites were selected based on sites permissive for HCV: (i) at the N-terminus of core, downstream of the first 12 amino acids (AAs) and upstream of a GSG linker, a P2A sequence to allow for cleavage, and the complete core protein (20); (ii) at the same position, but followed by rat ubiquitin downstream of P2A, allowing cleavage to generate the native N-terminus of core; (iii) between p7 and NS2, with Fluc followed by P2A and NS2 lacking the first AA (21); and (iv) between duplicated NS5A/NS5B cleavage sites (22) (**Figure 1A**). *In vitro* transcribed (IVT) RNA of these reporter constructs was transfected or electroporated into McA-RH7777.hi rat hepatoma cells with increased permissiveness to NrHV (9), with wildtype (WT) RHVcc-1 as a positive control. Cultures were followed for at least 4 weeks by antigen staining and luciferase measurements. Robust antigen staining or luciferase signal was never observed for NrHV-Fluc (i), (ii) and (iii). Only low luciferase signal was detectable, as exemplified for >40 days post-electroporation of NrHV-Fluc (i) (**Figure 1B**). For construct (iv), robust luciferase signal and viral RNA titres were observed early after transfection but these rapidly declined, comparable to a non-replicating pol(–) control (**Figure 1C**). To exclude that the observed deficiencies in viral replication were due to elements of the Fluc sequence, we additionally engineered constructs (iii) and (iv) with UpA/CpG-optimised enhanced green fluorescent protein (EGFP). However, no EGFP signal was observed by microscopy during >4 weeks of monitoring following transfection of IVT RNA. Thus, robust replication was not observed for any of the tested reporter constructs.

**Figure 1.**
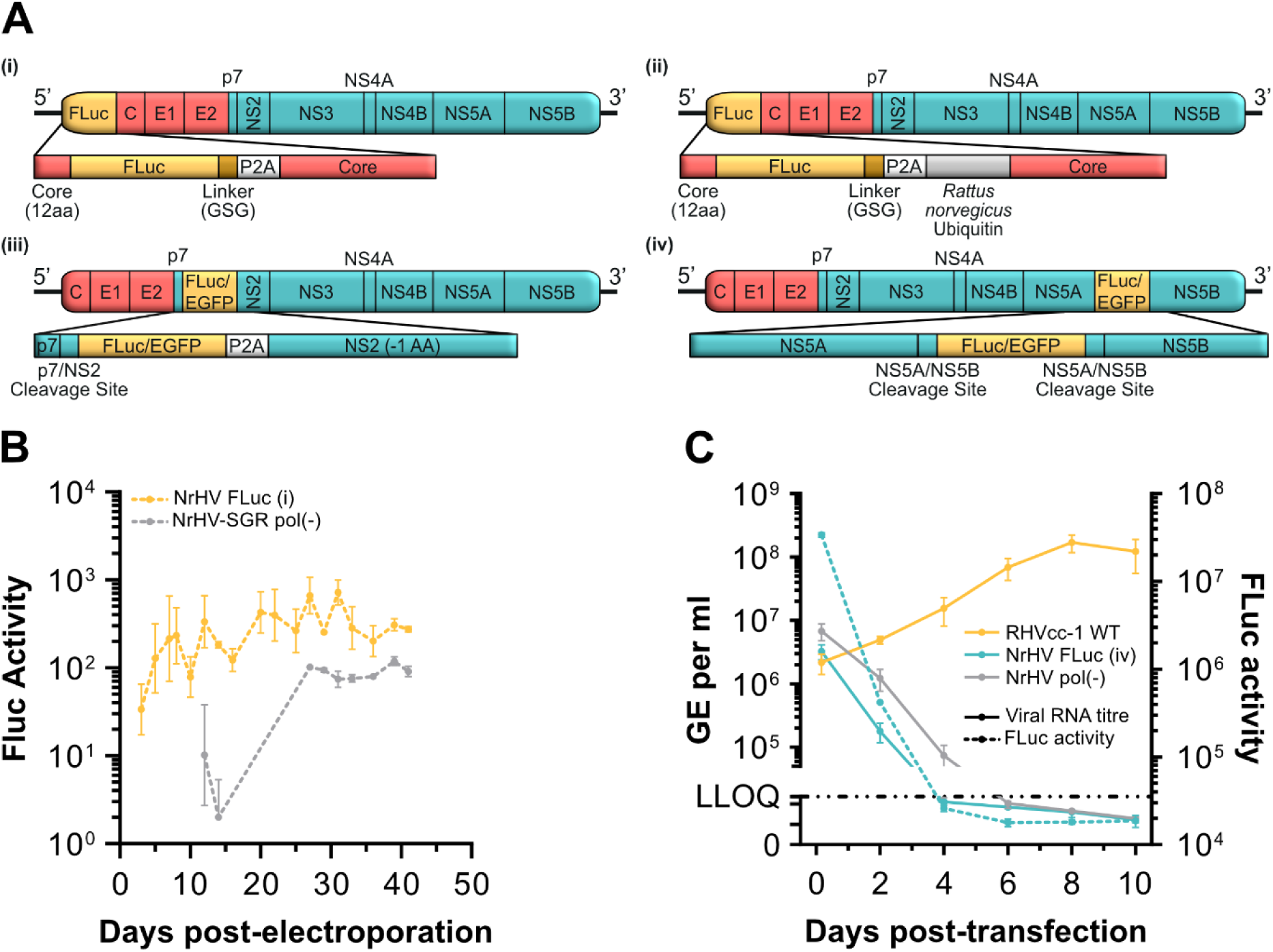
Low permissiveness for insertion of reporter genes into the NrHV ORF. **(A)** Schematic representations of four NrHV reporter constructs with insertions at the following sites: (i) FLuc inserted downstream of the first 12 amino acids (AA) of core and upstream of a GSG linker, a P2A sequence, and the full core protein; (ii) the same design as reporter (i) but with *Rattus norvegicus* ubiquitin inserted between the P2A sequence and core; (iii) either FLuc or EGFP inserted between p7 and NS2, downstream of the p7/NS2 cleavage site and upstream of a P2A sequence and NS2 lacking the first AA ; (iv) either FLuc or EGFP inserted between duplicated NS5A/NS5B cleavage sites. **(B)** FLuc activity for NrHV-Fluc (i) and a replication deficient NrHV sub-genomic FLuc replicon over time after electroporation of *in vitro* transcribed RNA into NrHV permissive McA-RH7777.hi rat hepatoma cells. Data points represent means ± standard deviation (SD) from triplicates for days 1-7, and duplicates thereafter. **(C)** Comparison of FLuc activity and RNA genome equivalents (GE) per mL of supernatant over time after transfection of *in vitro* transcribed RNA of RHVcc-1, reporter NrHV-Fluc (iv), or a replication deficient NrHV sub-genomic FLuc replicon into NrHV permissive McA-RH7777.hi rat hepatocytes. The lower limit of quantification (LLOQ) for RT-qPCR was 5x10^4^ GE/mL. Solid lines indicate RNA titres while dashed lines represent FLuc activity. Data points represent means ± SD from triplicates.

### Active replication of NrHV with a HiBiT tag inserted into NS2 or NS5A

Reasoning that full-length reporter genes may not be readily accommodated within the NrHV genome due to size restrictions, we instead inserted sequence encoding the 11 AA HiBiT tag at 4 positions in the NrHV ORF. The small HiBiT tag constitutes a functional NanoLuc luciferase enzyme when bound to the larger complementary LgBiT protein, allowing for quantification of HiBiT-tagged viral proteins upon lysis and addition of the LgBiT protein. Due to the requirement of appending the HiBiT tag onto proteins, the selected insertion sites differed but were again based on sites permissive for HCV: (i) after the first 2 AA of core, upstream of a GSSG linker and the remainder of core (23); (ii) after the first 3 AA of core, upstream of a GSSG linker, rat ubiquitin, and the full core protein; (iii) at the N-terminus of NS2, downstream of a duplication of the first 4 AAs of NS2 and upstream of a GSSG linker (24); and (iv) after AA 2322 within NS5A domain III, downstream of a GSSG linker (25, 26) (**Figure 2A**). Of note, conservation of NS5A domain III between NrHV and HCV, and even between HCV isolates, is low, challenging identification of the insertion site corresponding to that successfully employed for HCV (**Figure 2B**). IVT RNA of these NrHV reporter constructs, and the RHVcc-1 control, was then electroporated into McA-RH7777.hi cells. The infection was followed over time using antigen staining and measurement of NanoLuc luciferase (**Figure 2C-D**). No antigen-positive cells or luciferase signal was detected for the NrHV-HiBiT-Core and NrHV-HiBiT-Core-Ubi constructs. In contrast, NrHV-HiBiT-NS2 and NrHV-HiBiT-NS5A replication was detectable by both luciferase and antigen staining. NrHV-HiBiT-NS2 showed antigen positive cell counts comparable to the RHVcc-1 WT control, with a luciferase signal that followed this trend. NrHV-HiBiT-NS5A showed reduced antigen positive cells at early time points but reached WT levels by 7 days post-electroporation. The luciferase signal observed for NrHV-HiBiT-NS5A correlated with the quantification of antigen positive cells over time. Further, NrHV-HiBiT-NS5A had increasingly higher luciferase signal compared to NrHV-HiBiT-NS2, with the difference increasing from ∼7-fold to >21-fold higher signal from 2 to 18 days post-electroporation.

**Figure 2.**
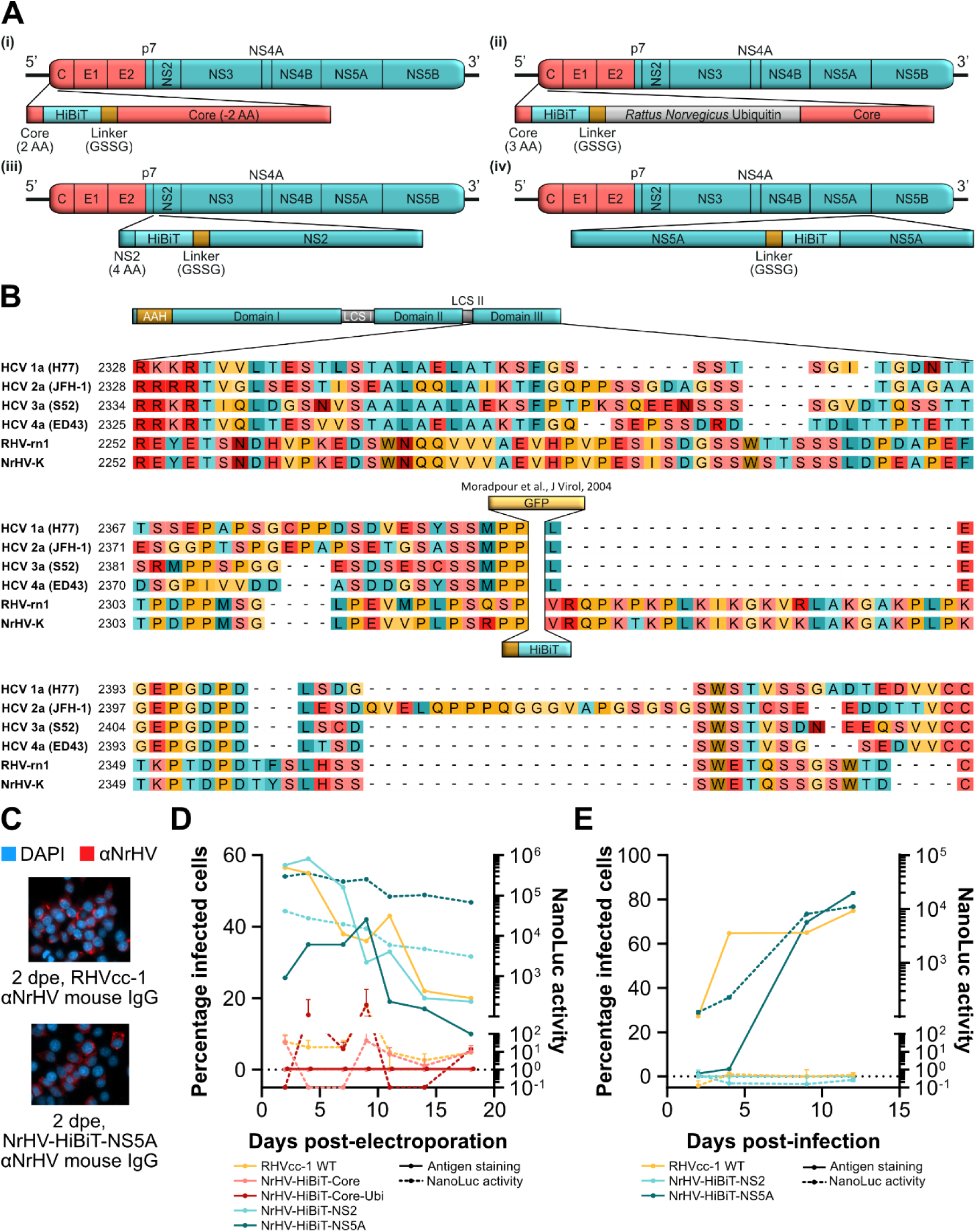
Insertion of the HiBiT tag into NS5A permits replication and infectious virus production. **(A)** Schematic representations of four NrHV reporter constructs with insertions of the HiBiT tag at the following sites: (i) NrHV-HiBiT-Core, at the N-terminus of core following the first two amino acids (AA) and upstream of a GSSG linker and core lacking the first 2 AA; (ii) NrHV-HiBiT-Core-Ubi, at the N-terminus of core following the first three AA and upstream of a GSSG linker, *Rattus norvegicus* ubiquitin, and the full core protein; (iii) NrHV-HiBiT-NS2, at the N-terminus of NS2 following the first 2 AA of NS2 and upstream of a GSSG linker and the full NS2 protein; and (iv) NrHV-HiBiT-NS5A, within NS5A domain III following AA 2322 and a GSSG linker. **(B)** MAFFT multiple sequence alignment of NS5A of HCV genotype 1a (H77, GenBank accession no. AF009606), HCV genotype 2a (JFH-1, GenBank accession no. AB047639), HCV genotype 3a (S52, GenBank accession no. GU814264), HCV genotype 4a (ED43, GenBank accession no. GU814265), RHV-rn1 (GenBank accession no. KX905133), and NrHV-K (GenBank accession no. PV553238) (43). Shown is NS5A domain III (based on domain definition for HCV) with indication of the insertion site of both the HiBiT tag and GFP in a functional HCV reporter (25, 26). Due to the low conservation of the depicted region, different alignment algorithms yielded variation in the resulting alignments. **(C-D)** Electroporation of IVT RNA of RHVcc-1 and HiBiT reporter constructs as shown in (A) into NrHV permissive McA-RH7777.hi rat hepatoma cells: (C) Imaging of antigen-stained RHVcc-1 and NrHV-HiBiT-NS5A 2 days post-electroporation (dpe), using total IgG from C57BL/6 mice that resolved NrHV infection, and (D) time course with solid lines indicating manually enumerated percentage infected cells and dashed lines representing NanoLuc activity. Data points represent individual values for percentage infected cells and means ± SD from duplicates for NanoLuc activity. **(E)** Infection of naïve McA-RH7777.hi rat hepatoma cells with supernatant collected from cells electroporated with RHVcc-1, NrHV-HiBiT-NS2, or NrHV-HiBiT-NS5A IVT RNA in the previous experiment (D), 9 days post-electroporation. Data points represent means ± SD from triplicates.

To assess the production of infectious particles, supernatants from NrHV-HiBiT-NS2 and NrHV-HiBiT-NS5A electroporated cells (9 days post-electroporation) were passaged onto naïve McA-RH7777.hi cells. No infection was observed for NrHV-HiBiT-NS2, while NrHV-HiBiT-NS5A exhibited a delayed spread of infection, eventually spreading to a fraction of cells comparable to the WT at 9 days post-infection (**Figure 2E**). The luciferase signal for NrHV-HiBiT-NS5A again closely followed quantification of percentage infected cells. In aggregate, this suggested NrHV-HiBiT-NS5A to be a functional reporter virus.

### Mutations in NS5A adapt NrHV-HiBiT-NS5A to cell culture

To determine if the increased percentage of infected cells and luciferase signal for NrHV-HiBiT-NS5A was due to the acquisition of adaptive mutations, we extracted viral RNA from supernatant collected 9 days post-infection (**Figure 2E**) and sequenced the viral ORF. This uncovered four non-synonymous mutations: R319K and V363L in E1 and W2289S and K2342N in NS5A (numbering according to the RHV-rn1 polyprotein). We therefore generated NrHV-HiBiT-KLSN containing all four mutations, and NrHV-HiBiT-SN containing only the two NS5A mutations (**Figure 3A**). IVT RNA of NrHV-HiBiT-SN and NrHV-HiBiT-KLSN was then electroporated into McA-RH7777.hi cells alongside the non-adapted NrHV-HiBiT-NS5A and WT RHVcc-1. NrHV-HiBiT-SN and NrHV-HiBiT-KLSN exhibited comparable percentage of antigen positive cells to WT RHVcc-1, while the non-adapted NrHV-HiBiT-NS5A was attenuated as previously observed (**Figure 3B**). After passage of supernatant from electroporated cells (11 days post-electroporation) to naïve McA-RH7777.hi cells, NrHV-HiBiT-SN and NrHV-HiBiT-KLSN exhibited similar infection kinetics to WT, suggesting that the two mutations in NS5A alone adapted NrHV-HiBiT-NS5A to cell culture (**Figure 3C**). For all further studies, NrHV-HiBiT-SN was therefore used to avoid the presence of additional mutations in the envelope proteins, which may impact studies of entry and neutralisation. To verify the stability of NrHV-HiBiT-SN, we electroporated its RNA into McA-RH7777.hi cells and serially passaged supernatant onto naïve cells four times each after 6-7 days. Sequencing of the viral ORF identified no coding mutations after 1-3 passages, and a single mutation, Q342P in E1, after 4 passages.

**Figure 3.**
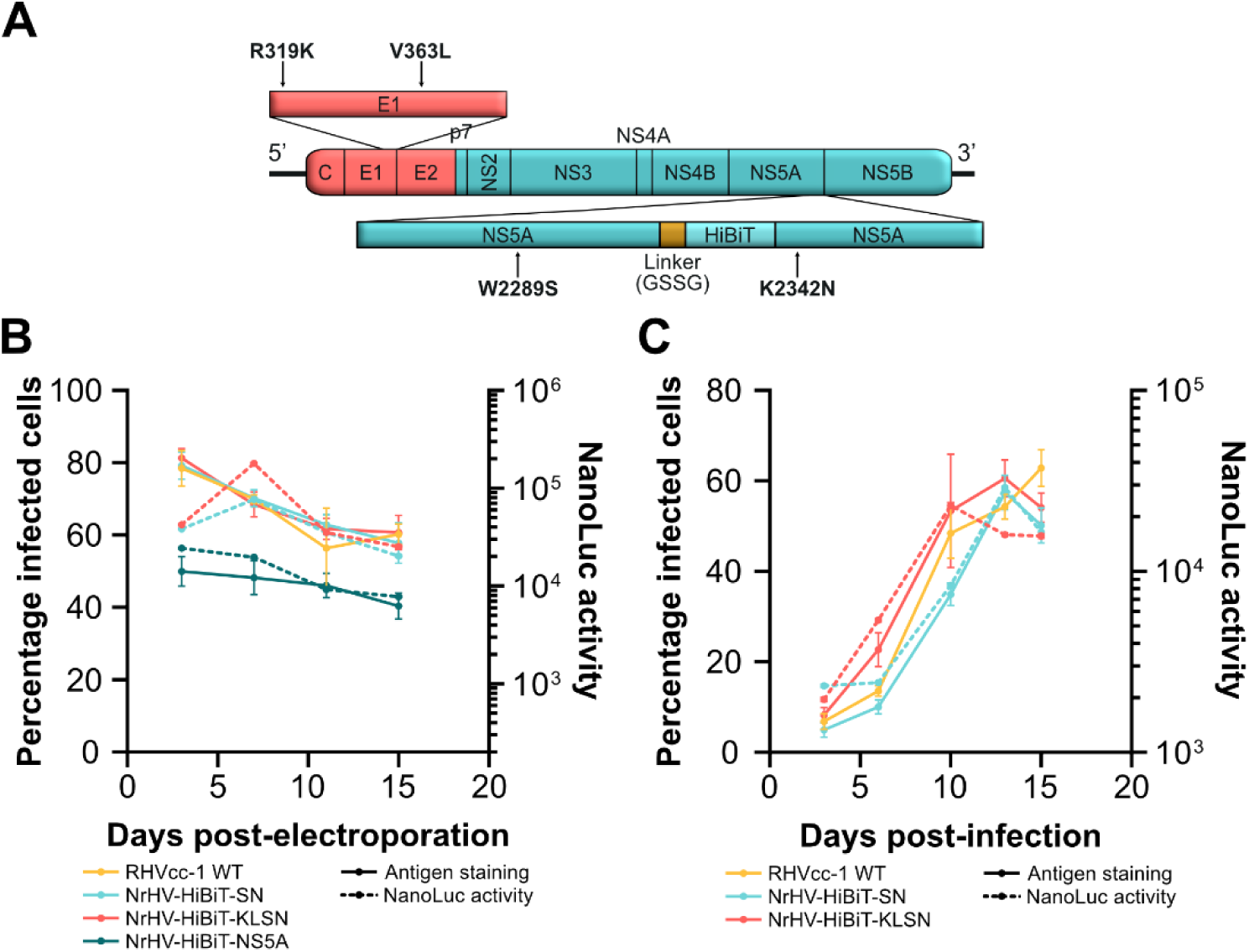
Adaptive mutations in NS5A promote fitness and stability of NrHV-HiBiT-NS5A *in vitro*. **(A)** Schematic representation of the location of the four adaptive mutations acquired by NrHV-HiBiT-NS5A. AA numbering for the labelled mutations corresponds to the RHV-rn1 polyprotein. **(B)** Electroporation of IVT RNA of RHVcc-1, NrHV-HiBiT-NS5A, NrHV-HiBiT-SN, and NrHV-HiBiT-KLSN, into McA-RH7777.hi rat hepatoma cells. Solid lines indicate manually enumerated percentage infected cells while dashed lines represent NanoLuc activity. **(C)** Infection of naïve McA-RH7777.hi rat hepatoma cells at MOI of 0.01 with supernatant collected from cells electroporated with RHVcc-1, NrHV-HiBiT-SN, or NrHV-HiBiT-KLSN IVT RNA in the previous experiment (B), 11 days post-electroporation. All data points represent means ± SD from triplicates.

### Viability of NrHV-HiBiT-SN *in vivo*

To permit luciferase-based readout of NrHV infection *in vivo*, we next inoculated four CB17-SCID mice each with 1 x 10^5^ genome equivalents (GE) of culture derived NrHV-HiBiT-SN or WT RHVcc-1. While titres of ∼10^7^-10^8^ GE/mL were detected for three of four WT RHVcc-1 inoculated mice throughout the course of the 21-day infection, NrHV RNA was not detectable in the sera of animals inoculated with NrHV-HiBiT-SN (**Figure 4A**).

**Figure 4.**
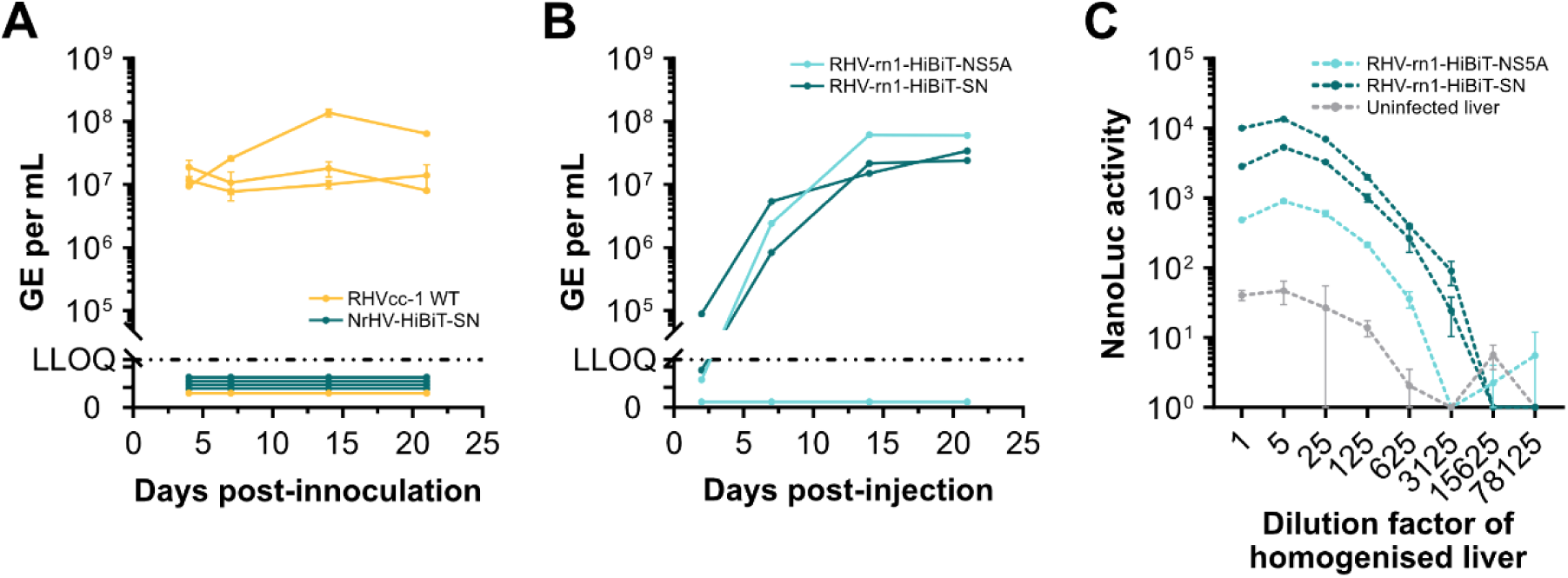
*In vivo* infection with NrHV HiBiT reporters. **(A)** Viral RNA titres in serum from CB17-SCID mice infected with 1x10^5^ GE of cell culture derived RHVcc-1 or NrHV-HiBiT-SN. data points represent means ± SD from duplicates. **(B)** Viral RNA titres in serum after intrahepatic injection of IVT RNA of RHV-rn1-HiBiT-NS5A or RHV-rn1-HiBiT-SN into Lewis rats. Data points represent means from duplicates. **(C)** NanoLuc luciferase activity after serial dilution of liver homogenates from Lewis rats taken on day 21 from the inoculation experiment shown in (B). Data points represent means ± SD from duplicates. The LLOQ for RT-qPCR was 5x10^4^ GE/mL.

Given that culture-adaptive mutations attenuate NrHV infection *in vivo* (11), we engineered the NS5A HiBiT tag into the original rat infectious RHV-rn1 clone, with or without the SN mutations (RHV-rn1-HiBiT-NS5A and RHV-rn1-HiBiT-SN). Viral RNA from these constructs were intrahepatically injected into two Lewis rats per construct and viral titres were followed for 4 weeks. Both animals inoculated with RHV-rn1-HiBiT-SN and one animal inoculated with RHV-rn1-HiBiT-NS5A had detectable titres by 7 days post-injection. NrHV RNA titres increased to >1x10^7^ GE/mL in all three animals 14 days post-injection (**Figure 4B**). Sanger sequencing confirmed that no changes to the HiBiT tag or engineered mutations were observed over the 4-week infection. Further, NanoLuc luciferase signal was detected from the homogenised livers of all NrHV positive rats (**Figure 4C**). The NanoLuc luciferase signal from the livers of both animals infected with RHV-rn1-HiBiT-SN was >10-fold higher than that of the animal infected with RHV-rn1-HiBiT-NS5A, suggesting the two mutations in NS5A also improved expression of the HiBiT tag *in vivo*.

### The NrHV HiBiT reporter allows high-throughput, accurate quantification of neutralising antibodies

To establish the correlation between NanoLuc luciferase signal and viral infectivity titres, McA-RH7777.hi cells were infected with increasing doses of NrHV-HiBiT-SN. Infection was quantified by manual counting of FFUs or by luciferase signal. The quantification correlated closely between the two methods across the dilution series, with luciferase exhibiting lower levels of variation at the highest dilutions (**Figure 5A**). To validate the utility of the NrHV-HiBiT-SN reporter for efficient quantification of nAbs, we used the previously described *in vitro* neutralisation assay (11, 17), and compared quantification by manual FFU counting and luciferase signal. Quantification of nAbs in serum from persistently infected rats was in high agreement between NanoLuc luciferase and manual FFU counts across the dilution series for the sera tested. Less variation was observed from quantification by luciferase (**Figure 5B**). Accordingly, NrHV-HiBiT-SN presents a high-throughput platform for the quantification of neutralising antibodies.

**Figure 5.**
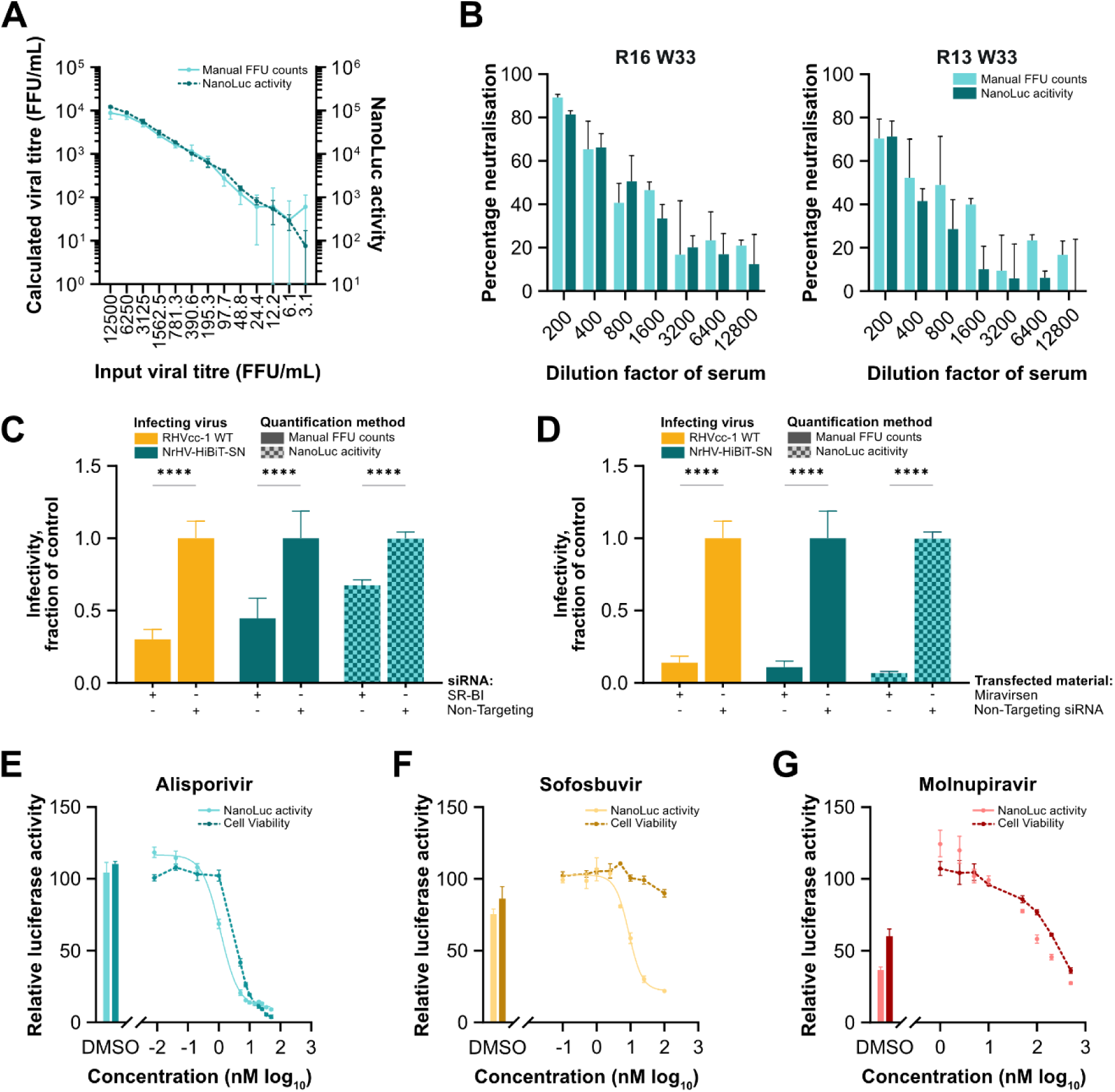
High-throughput quantification of neutralising antibodies, host factor inhibitors, and antiviral compounds using the NrHV-HiBiT-SN reporter. **(A)** Infection of McA-RH7777.hi rat hepatoma cells with decreasing titres of NrHV-HiBiT-SN. Viral infectivity titres were calculated through manual counts of FFUs after 48 hours and compared to NanoLuc luciferase signal. Data points represent means ± SD from triplicates. **(B)** NrHV-HiBiT-SN was exposed to dilutions of sera from chronically infected rats (R16 week 33, R13 week 33, (11)) before infection of McA-RH7777.hi cells. Percentage neutralisation compared to incubation with only culture medium was quantified through either manual counts of FFUs or NanoLuc luciferase. Data points represent means ± SD from triplicates. **(C-D)** Assessment of viral infection after transfection of siRNA or LNA to knock-down SR-BI (C) or inhibit miR-122 (D) relative to that from cells transfected with non-targeting siRNA. Bars of solid colour indicate manually enumerated FFUs while checkered bars represent NanoLuc activity. Data points represent means ± SD from eight replicates. **(E-G)** NanoLuc luciferase signal from McA-RH7777.hi cells electroporated with NrHV-HiBiT-SN treated with increasing doses of alisporivir (E), sofosbuvir (F), or molnupiravir (G). Values are given as percentage luciferase signal relative to untreated control wells. DMSO-only controls are shown for comparison. In addition, luciferase measurements of cell viability are given relative to untreated control wells using the CellTiter-Glo luminescent cell viability assay. Data points represent means ± SD from triplicates. Two-way ANOVA and Šídák’s multiple comparisons test were used to determine effects of siRNA or LNA transfection (C-D). Asterisks indicate p-values as follows: * p<=0.05, ** p<=0.01, *** p<=0.001, **** p<=0.0001.

### Efficient assessment of host factor interactions and antiviral compounds using the NrHV HiBiT reporter

To also demonstrate the utility of the HiBiT reporter in studies of interactions with host factors, we next focused on the entry factor SR-BI and the replication factor miR-122. McA-RH7777.hi cells were transfected with either SR-BI-targeting siRNA, non-targeting siRNA, or the miR-122 inhibitor miravirsen, prior to infection with WT RHVcc-1 or NrHV-HiBiT-SN. Quantification of NrHV-HiBiT-SN infection using luciferase again correlated well with the manual FFU counts of both WT RHVcc-1 and NrHV-HiBiT-SN infections for SR-BI siRNA and miravirsen-treated cells (**Figure 5C-D**). To next assess whether cyclophilin A (CypA), a cis-trans isomerase and host factor for HCV (27), also plays a role for NrHV, we electroporated McA-RH7777.hi cells with NrHV-HiBiT-SN prior to exposure to the CypA inhibitor, alisporivir (28). Although alisporivir had a clear dose-dependent effect on NrHV infection with an EC_50_ of 1.12 µM, the therapeutic window was narrow with apparent cytotoxicity at <10-fold higher doses (**Figure 5E**). Nonetheless, this suggested that CypA may act as a host factor for NrHV, similarly to HCV.

To similarly establish the utility of NrHV-HiBiT-SN to quantify activity of antiviral compounds, NrHV-HiBiT-SN was electroporated into McA-RH7777.hi cells and exposed to the HCV nucleotide/nucleoside inhibitors, sofosbuvir or molnupiravir (29, 30). A dose-dependent antiviral effect absent of cytotoxicity was observed for sofosbuvir against NrHV-HiBiT-SN (**Figure 5F**). Luciferase-derived results yielded an EC_50_ of 8.92 µM, correlating with the previously determined EC_50_ of 16 µM obtained for an NrHV sub-genomic replicon (NrHV-SGR) (9). Molnupiravir did not exhibit antiviral effect at non-cytotoxic concentrations (**Figure 5G**).

These findings highlight the utility of the NrHV-HiBiT-SN reporter as a tool for use in the characterisation of host-virus interactions and antiviral compounds for this hepacivirus model.

## Discussion

In this study, we generated an NrHV reporter virus for use in high-throughput analyses. While the insertion of full-length reporter genes proved to be highly attenuating, the insertion of the HiBiT tag (18) into NS5A domain III of the culture adapted RHVcc-1 (11) produced a reporter virus (NrHV-HiBiT-SN) with *in vitro* viability comparable to the parental clone after the acquisition of two mutations in NS5A. Engineering the HiBiT tag into NS5A of the original *in vivo* infectious RHV-rn1 clone (10, 13) further permitted productive infection in rats. The NrHV-HiBiT-SN reporter facilitated straightforward quantification, providing means to study viral replication, neutralising antibodies, and host factor interactions in a high-throughput manner. Using this reporter, we identified CypA as a potential novel host factor for NrHV. These findings expand our knowledge of NrHV as an HCV model and demonstrate the utility of this reporter in the further characterisation and study of NrHV.

Initial attempts to insert full-length FLuc or EGFP into culture adapted NrHV revealed key differences in the tolerance of insertions into the viral genome between HCV and NrHV. Despite each of the four insertion sites explored in this study having been inspired by functional HCV reporters (20–22), insertion of reporter genes at none of these sites was tolerated for NrHV replication. A transposon screen, as done for HCV and other viruses (31), could potentially aid in the discovery of acceptable insertion sites for NrHV and clarify the distinct differences between NrHV and HCV with regards to the tolerance of genomic modifications.

For HCV, the initial nucleotides of the ORF constitute part of domain IV of the IRES and are critical for efficient translation (32). While a similar structure has not been predicted for NrHV (13), this could explain why both HiBiT-Core reporters were replication incompetent, whereas an NrHV-SGR with FLuc insertions following the first 12 codons replicated efficiently (9). While the HiBiT insertions may have also altered the integrity and function of the Core protein itself, that would not be predicted to impact RNA replication. Experimental characterisation of the secondary structures of NrHV RNA would be needed to understand the role of the first nucleotides of the ORF in viral replication and translation.

The insertion of the HiBiT tag at the N-terminus of NS2 did not affect replication, consistent with observations that NS2 itself is not critical for HCV replication (33). It did, however, disrupt virus production, which is in accordance with the importance of NS2 for HCV virion morphogenesis and assembly (21, 34), with mutations in the first 5 AA residues heavily attenuating infectious virus production (35). This N-terminal insertion may therefore interfere with homologous functions for NrHV. The insertion of a HiBiT tag at this site was, however, permitted for HCV (24), alluding further to discrepancies in the tolerance to ORF modifications between the two viruses.

The NrHV-HiBiT-NS5A reporter was capable of both replication and infectious particle production *in vitro*. The insertion site in NS5A domain III was inspired by the insertion of GFP at the corresponding site in HCV NS5A (25, 26) and selected after alignment of NrHV NS5A with that of several HCV genotypes. The dissimilarity of NrHV and HCV in this region complicated the identification of corresponding sites, and the NrHV insertion may therefore not correspond exactly to that in HCV. Nonetheless, the large variability in this region, even among HCV genotypes (36), may suggest plasticity and flexibility for insertions. Of the four identified putative adaptive mutations for the NrHV-HiBiT-NS5A reporter, only the two mutations in NS5A were necessary to accommodate the HiBiT tag, with both NrHV-HiBiT-SN and NrHV-HiBiT-KLSN having fitness comparable to the parental RHVcc-1 virus *in vitro*. The two adaptive mutations in E1 have not been observed in previous studies, although V363L in E1 is in close proximity to several mutations involved in the adaptation of NrHV to mice (10). We speculate that the two E1 mutations were co-selected with little influence on viral fitness.

Unproductive infection of SCID mice with culture derived NrHV-HiBiT-SN was unexpected given successful infections with culture derived RHVcc-1 (11). This indicated that the combination of cell culture adaptive mutations and the insertion of the HiBiT tag may have attenuated the reporter virus beyond recovery. However, when re-engineered into the RHV-rn1 rat infectious clone (13), intrahepatic inoculation of both RHV-rn1-HiBiT-NS5A and RHV-rn1-HiBiT-SN RNA led to productive infection in rats. However, whereas RHV-rn1 infection or intrahepatic RNA inoculation typically leads to serum viremia of ∼10^9^ GE/mL in rats (11, 17), levels for the HiBiT reporters only reached 10^7^-10^8^ GE/mL. Long-term infection in rats may reveal further *in vivo* adaptive mutations to increase fitness. Nonetheless, the HiBiT sequence was stable in both reporters over the 4-week course of infection. Additionally, luciferase signal was observed from infected rat livers in a dose-dependent manner, confirming the integrity of the HiBiT tag. We observed ∼10-fold higher luciferase signal from the livers of animals infected with RHV-rn1-HiBiT-SN compared to RHV-rn1-HiBiT-NS5A, suggesting that the SN mutations also compensate for the insertion of the HiBiT tag *in vivo.* While this demonstrated the viability of the NrHV-HiBiT-SN reporter *in vivo*, direct luciferase monitoring would depend on co-expression of the complementary LgBiT protein, for example in a transgenic mouse model. Although the HiBiT system has been utilised for live imaging of viral infection *in vivo*, permitted by the prior injection of LgBiT expressing tumour cells (37), delivery of the LgBiT protein itself to the tissue of interest has proven difficult (38). Future development of a virus containing a full-length reporter gene would therefore still be of interest for utilisation of live imaging *in vivo*.

NrHV infection was reliably and more stably measured using the NrHV-HiBiT-SN reporter compared to manual FFU counts. Accordingly, the NrHV reporter provides a high-throughput method for the assessment of the neutralising capability of antibodies, a tool that can accelerate ongoing vaccine studies using the NrHV model. More broadly, we demonstrate the benefit of this reporter in the study of viral host factors, with both luciferase signal and manual FFU counts reporting comparable attenuation of infection during knock-down or depletion of the known NrHV host factors SR-BI and miR-122 (9–11). The lack of additional envelope mutations avoids further issues of interpreting data, e.g. from studies of neutralisation or viral entry, and the automated process of luciferase measurement reduces the influence of human error and bias on data collection. CypA is a critical host factor for HCV, providing a target for host-targeting therapeutic approaches (39). Our data here indicated that CypA may similarly be a host factor for NrHV, although only a narrow dose window demonstrated antiviral effects of alisporivir absent of cytotoxicity. The sensitivity of infections to alisporivir also differed, with NrHV-HiBiT-SN having an EC_50_ of 1.12 µM compared to 0.03 µM for HCV (40). Although this could be indicative of diverging reliance on CypA, it could also be the effect of lower efficacy of alisporivir mediated inhibition of rat compared to human CypA, or reflect differences in cellular uptake. This finding nonetheless corroborates the use of this reporter in the study of host-virus interactions. Finally, luciferase measurements revealed an EC_50_ of 8.92 µM for the inhibition of NrHV by sofosbuvir, corresponding to previous observations from the NrHV sub-genomic replicon (9). For molnupiravir, we did not observe antiviral effects at non-cytotoxic concentrations.

In summary, we generated a stable NS5A-HiBiT-tagged NrHV reporter virus permitting high-throughput studies of viral infection, yielding results in concordance with previous manual methods. This work thereby provides a flexible high-throughput tool for the NrHV model, facilitating the study of host factors, antivirals, and neutralising antibodies. NrHV holds great promise as a model for HCV, permitting the study of intrahepatic pathology in an immunocompetent setting as well as the assessment of vaccine platforms and candidates in a rodent challenge model. Future work to similarly introduce the HiBiT tag into other NrHV strains (16), would further allow high-throughput assessment of nAb cross-reactivity, furthering work towards a multivalent vaccine. Knowledge from the NrHV model on the best performing vaccine candidates, or combinations thereof, could then be applied with greater confidence to trials in high-incidence HCV patient cohorts (41). This high-throughput reporter will ease further characterisation of NrHV as a model and accelerate vaccine studies, permitting increased efforts in the critical quest for an HCV vaccine.

## Materials and methods

### Animal experiments

Animal experiments were done at the Department of Experimental Medicine, University of Copenhagen, under protocol 2021-15-0201-01063 and 2022-15-0201-01292 approved by the national Animal Experiments Inspectorate. All work was consistent with an affirmative response to the ARRIVE 10 questionnaire. For infection experiments, female CB17-SCID (CB-17/Icr-Prkdcscid/scid/Rj) mice were inoculated intraperitoneally with 1x10^5^ GE of cell culture derived NrHV under isoflurane anaesthesia. RNA launch in rats was conducted in female LEWOrl/Rj rats (Janvier) by percutaneous intrahepatic injection under isoflurane anaesthesia of 10 µg IVT NrHV genomic RNA dissolved in 100 µL sterile PBS distributed in two hepatic locations, each receiving 5 µg. Mice were bled via facial vein puncture, while rat blood was sampled by tail vein puncture under isoflurane anaesthesia according to guidelines for blood sampling. All animals had access to food (SAFE D03, SAFE Complete Care Competence, Rosenheim, Germany) and water *ad libitum* and were housed by gender (except for breeders) in Innovive IVC caging containing wood chip bedding, shelters, nesting material, and biting sticks on a 12-hour light dark cycle. All experimentation was conducted during light cycle.

### Cells and antibodies

McA-RH7777.hi rat hepatoma cells permissive to infection by NrHV (9) were maintained in DMEM (Thermo Fisher Scientific) supplemented with 10% FBS, 100 U/mL penicillin, and 100 µg/mL streptomycin (Pen Strep: Sigma-Aldrich) at 37°C, 5% CO_2_. Cells were passaged every 2-3 days using trypsin-EDTA (Sigma-Aldrich).

Polyclonal NrHV antibody was obtained by purification of total IgG from sera collected from C57BL/6 mice 6 weeks post-infection with mouse-adapted NrHV (15).

### Cloning of NrHV constructs

All NrHV reporter constructs were generated through the ligation of single or multiple amplicons using the In-Fusion Snap Assembly Master Mix (Takara Bio), following manufacturer’s instructions, or through megaprimer mutagenesis unless otherwise stated. All megaprimer fragments were gel purified using the Zymoclean Gel DNA Recovery Kit (Zymo Research). Megaprimer PCR amplifications were completed using 200 ng of purified megaprimer fragment and 50 ng of template plasmid with consistent thermocycler conditions (1 cycle of 98°C for 30 s; 20 cycles of 98°C for 10 s, 48°C for 1 minute, 72°C for 20 minutes; 1 cycle of 72°C for 20 minutes; hold at 4°C). All PCR reactions were carried out using Q5 Hot Start High-Fidelity 2X Master Mix (New England Biolabs) unless using a QuikChange Site-Directed Mutagenesis Kit (Agilent Technologies), for which the proprietary polymerase was used. All clones were sequence verified over the entirety of the viral genome using Sanger sequencing (Macrogen).

To engineer an RHVcc-1 reporter with CpG/UpA optimised FLuc (19) inserted at the N-terminus of core, primer pair TS-O-01253/TS-O-01254 (**Table A1**) was used to amplify the 5’ UTR and FLuc insert from a monocistronic NrHV-SGR (9), including a GSG linker and P2A sequence as a 3’ overhang. The remaining sequence of the cell culture adapted RHV-rn1 plasmid, pRHVcc-1 (11), was amplified with the primer pair TS-O-02155/TS-O-01256 and the two fragments were ligated. The resulting plasmid contained a frameshift mutation that was corrected using a megaprimer generated from primer pair TS-O-01283/TS-O-00260. To insert *Rattus norvegicus* ubiquitin downstream of the P2A sequence, this frameshift-corrected reporter was linearised using the primer pair TS-O-01425/TS-O-01426 and ligated with *Rattus norvegicus* ubiquitin synthesised by IDT (IDT_Ubi_BuildingBlock, **Table A1**).

Insertion of CpG/UpA optimised FLuc or EGFP between p7 and NS2 was achieved through ligation of linearised RHVcc-1 sequence and plasmid backbone, amplified using the primer pair TS-O-01221/TS-O-01222, and CpG/UpA optimised FLuc or EGFP amplified from NrHV sub-genomic replicons containing the appropriate reporter gene (9) using primer pairs TS-O-01223/TS-O-01224 or TS-O-01227/TS-O-01228.

To insert CpG/UpA optimised FLuc in between duplicated NS5A/NS5B cleavage sites, the sequence and plasmid backbone of an RHV-rn1 construct containing the S1757A and T2373A cell culture-adaptive mutations (11) and the FLuc reporter gene from a RHV-rn1-SGR construct (9), containing a fragment of the NS5A/NS5B cleavage site at the 5’ and 3’ ends, were amplified using the primer pairs TS-O-00889/TS-O-00890 and TS-O-01098/TS-O-01099 respectively, and were subsequently ligated. CpG/UpA optimised EGFP was inserted at the same site using an 800 bp megaprimer amplified from an RHV-rn1-SGR-EGFP construct (9) using the primer pair TS-O-00305/TS-O-00306, similarly containing the NS5A/NS5B cleavage site at both 5’ and 3’ ends. Missing cell culture adaptive mutations were introduced to both reporter constructs using a 5387 bp megaprimer, including the L586S mutation, amplified from the same RHV-rn1 construct containing the S1757A and T2373A cell culture-adaptive mutations (11) using primer pair TS-O-01105/TS-O-01106. The T2373A mutation was engineered into the 2^nd^ NS5A/NS5B cleavage site using the QuikChange Lightning Site-Directed Mutagenesis Kit (Agilent Technologies) following manufacturer’s instructions with the primer pair TS-O-01184/TS-O-01185, thereby placing this adaptive mutation in both repeats of the cleavage site.

To insert the HiBiT tag into RHVcc-1, we ligated amplicons of the entirety of the RHVcc-1 genome and plasmid backbone amplified by the primer pairs TS-O-01591/TS-O-01592, TS-O-01597/TS-O-01598, and TS-O-01599/TS-O-01600 containing the HiBiT tag and linker as overhangs at the 5’ and 3’ ends. Primer pair TS-O-01595/TS-O-01596 enabled amplification of *Rattus norvegicus* ubiquitin, thus ligation with the full-length RHVcc-1 amplicon produced by the primer pair TS-O-01593/TS-O-01594 permitted the insertion of ubiquitin downstream of the HiBiT tag and GSSG linker.

To generate the NrHV-HiBiT-SN and NrHV-HiBiT-KLSN mutants of NrHV-HiBiT-NS5A, the primer pairs TS-O-01713/TS-O-01714 and TS-O-01711/TS-O-01712 were used to create amplicons containing the desired mutations within E1 and NS5A, respectively. These amplicons were used for megaprimer amplification, allowing mutation first of the two positions in NS5A and subsequently the two additional positions in E1.

The RHV-rn1-HiBiT clone was produced through ligation of an amplicon of the entirety of the RHV-rn1 plasmid produced by the primer pair TS-O-01599/TS-O-01600, introducing the HiBiT tag and GSSG linker to NS5A domain III.

To produce the RHV-rn1-HiBiT-SN mutant, HiBiT-adaptive mutations were introduced to RHV-rn1-HiBiT using two methods. The W2289S mutation in NS5A was engineered using the megaprimer method described above, and the K2342N mutation was added using QuikChange XL Site-Directed Mutagenesis Kit (Agilent Technologies) following manufacturer’s instructions with the primer pair TS-O-02128/TS-O-02129.

### RNA *in vitro* transcription, transfection and electroporation

Viral RNA transcripts for electroporation were generated from 2.5 µg of *MluI*-linearised plasmid using the T7 RiboMAX express large scale RNA production system (Promega), before being DNase treated with RQ1 RNase-free DNase (Promega) for 30 minutes on ice. IVT generated RNA was then purified using RNA Clean & Concentrator-25 (Zymo Research) prior to storage at -70°C.

Prior to transfections, 300,000 McA-RH7777.hi cells (9) were seeded in 6-well plates pre-coated with 0.1% gelatine. To transfect IVT RNA, Lipofectamine 2000 transfection reagent (Thermo Fisher Scientific) was diluted in Opti-MEM (Thermo Fisher Scientific) at a ratio of 1:50. For each construct to be transfected, 5 µg IVT RNA was then diluted in 250 µL Opti-MEM. Diluted Lipofectamine 2000 transfection reagent and IVT RNA were then mixed at a ratio of 1:1 and incubated at room temperature for 20 minutes. Growth media was then removed from wells containing the previously seeded McA-RH7777.hi cells. Cells were washed with phosphate buffered saline (PBS) and 1.5 mL Opti-MEM was added before 500 µL of the transfection mixture was gently pipetted onto cells. Plates were then incubated at 37°C, 5% CO_2_.

For electroporation, McA-RH7777.hi cells (9) were washed twice with ice-cold PBS before being resuspended at a cell density of 1.5 x 10^7^ cells/mL in either PBS supplemented with 1:50 ATP and 1:50 glutathione or cytomix (120 mM KC1; 0.15 mM CaCl_2_; 10 mM K_2_HPO_4_/KH_2_PO_4_, pH 7.6; 25 mM HEPES, pH 7.6; 2 mM EGTA, pH 7.6; 5mM MgCl2, pH adjusted with KOH; 2mM ATP and 5mM glutathione). Once diluted, 400 µL of cell suspension was added to 5 µg of *in vitro* transcribed RNA and transferred to a 4 mm electroporation cuvette (Bio-Rad). The following parameters on a Bio-Rad Gene Pulser xCell electroporation system were used for exponential decay electroporation: 270 V, 975 µF, ∞ ohm. Electroporated cells were immediately resuspended in DMEM pre-warmed to 37°C and subsequently transferred to 15 cm culture dishes coated with filtered 0.1% gelatine, prepared from porcine gelatine powder (Sigma-Aldrich) dissolved in MilliQ water. Post-electroporation, cells were split and supplied with fresh media every 2 days.

### Transfection of IVT RNA, siRNA, and LNA

To transfect McA-RH7777.hi cells (9) with siRNA or miravirsen (42), 300,000 cells were seeded in 6-well plates pre-coated with 0.1% gelatine. Lipofectamine RNAiMAX transfection reagent (Thermo Fisher Scientific) was then diluted in Opti-MEM (Thermo Fisher Scientific) at a ratio of 1:16.67. SR-BI siRNA (ON-TARGETplus SMARTpool, Horizon Discovery LTD), non-targeting siRNA pool (ON-TARGETplus Non-targeting siRNA #4) or miravirsen (+CC*+AT*T*+G+TC*A*+CA*+CT*+C+C; + designating LNA bases and * phosphorothioate DNA backbone; Qiagen) were reconstituted in RNase-free water and diluted in Opti-MEM to the desired concentration for transfection. Diluted siRNAs and LNAs were mixed with diluted Lipofectamine RNAiMAX transfection reagent at a ratio of 1:1 and left to incubate at room temperature for 5 minutes. Onto each well containing seeded cells covered in 1.7 mL DMEM supplemented with 3% FBS, 100 U/mL penicillin, and 100 µg/mL streptomycin (Pen Strep: Sigma-Aldrich), 300 µL of this transfection mixture was carefully pipetted prior to incubation for 4-6 hours at 37°C, 5% CO_2_. Transfection of siRNA and LNA was carried out twice, 2 days and 1 day prior to infection with NrHV constructs.

### Extraction, quantification, and sequencing of viral RNA

To extract viral RNA from supernatant of infected cell cultures, 250 µL of supernatant was mixed with 750 µL TRIzol LS reagent (Thermo Fisher Scientific) and incubated for 5 minutes at room temperature. For extraction of viral RNA from rodent serum samples, 25 µL of sera was diluted in 225 µL chilled PBS prior to mixing with TRIzol LS reagent. To this mixture, 200 µL of chloroform was added before being shaken vigorously for 15 s and left at room temperature for a further 3 minutes before being centrifuged at 12,000 g for 15 minutes at 4°C. After centrifugation, the aqueous phase was recovered and mixed with 450 µL of anhydrous ethanol and transferred to an RNA Clean & Concentrator-5 column (Zymo Research), with RNA purification and concentration being carried out following manufacturer’s instructions. Detection and quantification of NrHV RNA was achieved through RT-qPCR with TaqMan Fast Virus 1-Step Master Mix (Thermo Fisher Scientific) using a LightCycler 480 thermocycler running the following protocol: 50°C for 30 minutes, 95°C for 5 minutes, followed by 40 cycles of 95°C for 15 s, 56°C for 30 s, and 60°C for 45 s, and finally 40°C for 10 s. Amplification was achieved using the primers TS-O-00561 and TS-O-00562 and probe TS-O-00563 (**Table A1**).

For whole ORF sequencing, following RNase inhibition by incubation with RNasin Plus RNase inhibitor (Promega) for 2 minutes at 65°C, cDNA of the viral ORF was generated using Maxima Minus H Reverse Transcriptase (Thermo Fisher Scientific) with TS-O-00319 as a 3’ cDNA primer, being incubated for 120 minutes at 50°C followed by 5 minutes at 85°C. RNA was degraded through a 20 minute incubation at 37°C with RNase H and RNase T1 (Thermo Fisher Scientific). The viral ORF was amplified using Q5 Hot Start High-Fidelity DNA Polymerase (New England Biolabs) with the primers TS-O-00316 and TS-O-00318 and the following thermocycler settings: 98°C for 30 s, 37 cycles of 98°C for 10 s, 65°C for 10 s, 72°C for 8 minutes, and a final cycle of 72°C for 10 minutes before being held at 4°C. Amplified ORFs were sent for Sanger sequencing (Macrogen).

### Detection of luminescence and fluorescence

To measure FLuc activity, cells were washed once with PBS after the removal of growth media and then lysed with passive lysis buffer. Luciferase was then measured with the luciferase assay system (Promega) following manufacturer’s instructions using a CLARIOstar luminometer (BMG Labtech).

To inspect EGFP signal, cells were seeded in 8-chamber slides, replacing DMEM (Thermo Fisher Scientific) with FluoroBrite DMEM (Thermo Fisher Scientific) once cells had settled to reduce autofluorescence. Chamber slides were manually inspected for fluorescent cells using a Carl Zeiss Axio Vert. A1 microscope.

For measurement of NanoLuc luciferase activity using the HiBiT Nano-Glo detection system (Promega), Nano-Glo Lytic Reagent was prepared following manufacturer’s instructions. Where luciferase was to be detected from cells seeded in 96-well plates, cells were seeded in opaque white 96-well cell culture plates coated with 0.1% gelatine, with a media volume of 50 µL. For detection of luciferase from pelleted cells, cells were first resuspended in PBS to the cell concentration at which they were pelleted, with 50 µL of the resuspension being added to an opaque white 96-well luciferase plate. An equal volume of Nano-Glo Lytic Reagent was added to samples and mixed by pipetting. Plates were placed on an orbital shaker for at least 10 minutes prior to detection and quantification of luminescence using a CLARIOstar luminometer (BMG Labtech).

To measure NanoLuc luciferase signal from infected Lewis rat livers, liver sections weighing ∼100 mg after trimming of fat and connective tissue were transferred to a 1.5 mL screw cap tube and mechanically homogenised in 500 µL PBS using a 1.5 mL disposable pestle. Homogenates were then centrifuged for 3 minutes at 100 g, 4°C, before being diluted as indicated. Dilutions were added to an opaque white 96-well luciferase plate and processed as above. Luciferase measurements were adjusted to control for variation in weight between liver samples.

### Infectivity titration

Prior to infectivity titration, 96-well culture plates were coated by a 2-hour incubation with 50 µL of 10 µg/mL laminin. Plates were washed 3 times with PBS to remove any unbound laminin before seeding 13,500 McA-RH7777.hi rat hepatoma cells (9) per well. Supernatants from infected cell cultures were thawed and diluted to create a 10-fold dilution series from 1:2 to 1:2000. Each dilution of virus stock was applied to the cells in triplicate and cells were incubated at 37°C, 5% CO_2_, for 48 hours after which antigen staining of NrHV was carried out with NrHV-specific mouse IgG and Alexa Fluor 594 goat-anti-mouse IgG as previously described (11, 15, 17). Images were captured at 50x magnification using a Carl Zeiss Axio Vert. A1 microscope. The FFU/mL titre was enumerated manually, with a single FFU being defined as a group of more than one infected cell with a distance of at least two uninfected cells from another FFU.

### Neutralisation assay

To determine the neutralising capacity of NrHV specific antibodies in serum from infected animals, 13,500 McA-RH7777.hi cells (9) were seeded in laminin-coated 96-well cell culture plates. Serum from infected animals was heat inactivated at 56°C for 30 minutes prior to use. Serum was then diluted 1:100 in DMEM supplemented with 10% FBS, 100 U/mL penicillin, and 100 µg/mL streptomycin (Pen Strep: Sigma-Aldrich), with this dilution being further serially diluted 1:2 to acquire dilutions ranging from 1:100 – 1:6400. Virus and dilutions of sera were mixed in an empty 96-well plate at a ratio of 1:1 and incubated for 1 hour at 37°C, 5% CO_2_. For positive controls, virus was mixed with DMEM at a ratio of 1:1 and for negative controls, only DMEM was used. After 1 hour, the serum/virus mix and controls were transferred onto the seeded McA-RH7777.hi cells and incubated at 37°C, 5% CO_2_ for 4 hours. Cells were then washed twice with PBS, covered with 100 µL DMEM and incubated for a further 44 hours. After this time, infections were either processed for NanoLuc luciferase or visualised and FFUs counted as described for infectivity titration. Percentage neutralisation was calculated through comparison of NanoLuc luciferase signal or FFU counts to the mean NanoLuc luciferase signal or FFU count of the control in triplicates.

### Treatment with DAAs and inhibitors

To evaluate the quantification of cyclophilin A inhibition and the effect of sofosbuvir and molnupiravir on NrHV by NanoLuc luciferase, IVT RNA of NrHV-HiBiT-SN was electroporated into McA-RH7777.hi cells. Following electroporation, 7,500 cells were seeded per well into a laminin-coated 96-well plate. Alisporivir was serially diluted to concentrations ranging from 50 µM-8 nM, while sofosbuvir and molnupiravir were diluted to concentrations ranging from 100 µM-0.1 µM and 500 µM-1 µM respectively. DMSO-only controls were prepared by mimicking the preparation of the lowest dilution of alisporivir, sofosbuvir, and molnupiravir, adding only DMSO. All dilutions were prepared in DMEM supplemented with 3% FBS. The diluted compounds were added in triplicate 24 hours post-electroporation. Cells were incubated at 37°C, 5% CO_2_, for 48 hours, after which fresh dilutions of the compounds were added and incubation continued for an additional 48 hours. Infections were then quantified using NanoLuc luciferase.

To assess the effect of these compounds on cell viability, McA-RH7777.hi cells were electroporated and treated with compounds as described above, with 7,500 cells seeded per well into a laminin-coated white opaque 96-well cell culture plate. Cell viability was then assessed using the CellTiter-Glo luminescent cell viability assay (Promega) following manufacturer’s instructions and using a using a CLARIOstar luminometer (BMG Labtech).

## Statistical analyses and multiple alignment

Multiple alignment of the protein sequence of NS5A of HCV genotype 1a (H77, GenBank accession no. AF009606), HCV genotype 2a (JFH-1, GenBank accession no. AB047639), HCV genotype 3a (S52, GenBank accession no. GU814264), HCV genotype 4a (ED43, GenBank accession no. GU814265), RHV-rn1 (GenBank accession no. KX905133), and NrHV-K (GenBank accession no. PV553238) was generated using the MAFFT version 7 web server using default settings, L-INS-i strategy, and the BLOSUM45 scoring matrix (43). Two-way ANOVA, Šídák’s multiple comparisons tests, and non-linear regression analyses were performed using GraphPad Prism 10.5.0.

## Acknowledgments

We acknowledge the Department of Infectious Diseases, Copenhagen University Hospital, Hvidovre and the Department of Immunology and Microbiology, University of Copenhagen for support, the Department of Clinical Microbiology and the Department of Pathology, Hvidovre Hospital, for sequencing resources, and the Department of Experimental Medicine, University of Copenhagen, for rodent husbandry and care. The present study was supported by the Independent Research Fund Denmark (1030-00426 to T.K.H.S), the Novo Nordisk Foundation (NNF19OC0054518, NNF19OC0055462 and NNF23OC0087091 to J.B.), and the Danish Cancer Society (R374-A22411 to T.K.H.S.). R.W. was supported by an Early Postdoc.Mobility Fellowship (P2BEP3_178527) and a Postdoc.Mobility Fellowship (P400PB_183952) from the Swiss National Science Foundation.

## References

1. World Health Organization. 2024. Global hepatitis report 2024: action for access in low-and middle-income countries. World Health Organization, Genève, Switzerland.

2. Manns MP, Maasoumy B. 2022. Breakthroughs in hepatitis C research: from discovery to cure. Nat Rev Gastroenterol Hepatol 19:533–550.

3. Bartenschlager R, Baumert TF, Bukh J, Houghton M, Lemon SM, Lindenbach BD, Lohmann V, Moradpour D, Pietschmann T, Rice CM, Thimme R, Wakita T. 2018. Critical challenges and emerging opportunities in hepatitis C virus research in an era of potent antiviral therapy: Considerations for scientists and funding agencies. Virus Res 248:53–62.

4. Offersgaard A, Bukh J, Gottwein JM. 2023. Toward a vaccine against hepatitis C virus. Science 380:37–38.

5. Burm R, Collignon L, Mesalam AA, Meuleman P. 2018. Animal Models to Study Hepatitis C Virus Infection. Front Immunol 9:1032.

6. Bukh J. 2012. Animal models for the study of hepatitis C virus infection and related liver disease. Gastroenterology 142:1279–1287.e3.

7. Kennedy MJ, Fernbach S, Scheel TKH. 2024. Animal hepacivirus models for hepatitis C virus immune responses and pathology. J Hepatol 81:184–186.

8. Berggren KA, Suzuki S, Ploss A. 2020. Animal Models Used in Hepatitis C Virus Research. Int J Mol Sci 21:3869.

9. Wolfisberg R, Holmbeck K, Nielsen L, Kapoor A, Rice CM, Bukh J, Scheel TKH. 2019. Replicons of a Rodent Hepatitis C Model Virus Permit Selection of Highly Permissive Cells. J Virol 93:e00733–19.

10. Billerbeck E, Wolfisberg R, Fahnøe U, Xiao JW, Quirk C, Luna JM, Cullen JM, Hartlage AS, Chiriboga L, Ghoshal K, Lipkin WI, Bukh J, Scheel TKH, Kapoor A, Rice CM. 2017. Mouse models of acute and chronic hepacivirus infection. Science 357:204–208.

11. Wolfisberg R, Thorselius CE, Salinas E, Elrod E, Trivedi S, Nielsen L, Fahnøe U, Kapoor A, Grakoui A, Rice CM, Bukh J, Holmbeck K, Scheel TKH. 2022. Neutralization and receptor use of infectious culture-derived rat hepacivirus as a model for HCV. Hepatology 76:1506–1519.

12. Tanaka T, Akaike Y, Kasai H, Yamashita A, Matsuura Y, Moriishi K. 2025. Identification of claudin-3 as an entry factor for rat hepacivirus. Proc Natl Acad Sci U S A 122:e2508736122.

13. Trivedi S, Murthy S, Sharma H, Hartlage AS, Kumar A, Gadi SV, Simmonds P, Chauhan LV, Scheel TKH, Billerbeck E, Burbelo PD, Rice CM, Lipkin WI, Vandegrift K, Cullen JM, Kapoor A. 2018. Viral persistence, liver disease, and host response in a hepatitis C-like virus rat model. Hepatology 68:435–448.

14. Hartlage AS, Murthy S, Kumar A, Trivedi S, Dravid P, Sharma H, Walker CM, Kapoor A. 2019. Vaccination to prevent T cell subversion can protect against persistent hepacivirus infection. Nat Commun 10:1113.

15. Wolfisberg R, Holmbeck K, Billerbeck E, Thorselius CE, Batista MN, Fahnøe U, Lundsgaard EA, Kennedy MJ, Nielsen L, Rice CM, Bukh J, Scheel TKH. 2023. Molecular Determinants of Mouse Adaptation of Rat Hepacivirus. J Virol 97:e0181222.

16. Thorselius CE, Kok A, Wolfisberg R, Fahnøe U, Kennedy MJ, Lundsgaard EA, Mikkelsen L, Larsen MK, Murthy S, Trivedi S, Kapoor A, Scheel TKH, Holmbeck K, Bukh J. 2025. In vivo and in vitro recombinant systems of a novel variant demonstrate cross-reactive neutralization for the HCV model virus, Norway rat hepacivirus. PLoS Pathog 21:e1013127.

17. Thorselius CE, Wolfisberg R, Fahnøe U, Scheel TKH, Holmbeck K, Bukh J. 2025. Norway rat hepacivirus resembles hepatitis C virus in terms of intra-host evolution and escape from neutralizing antibodies. J Hepatol 83:870–880.

18. Schwinn MK, Machleidt T, Zimmerman K, Eggers CT, Dixon AS, Hurst R, Hall MP, Encell LP, Binkowski BF, Wood KV. 2018. CRISPR-mediated tagging of endogenous proteins with a luminescent peptide. ACS Chem Biol 13:467–474.

19. Witteveldt J, Martin-Gans M, Simmonds P. 2016. Enhancement of the replication of hepatitis C virus replicons of genotypes 1 to 4 by manipulation of CpG and UpA dinucleotide frequencies and use of cell lines expressing SECL14L2 for antiviral resistance testing. Antimicrob Agents Chemother 60:2981–2992.

20. Zhang Y, Weady P, Duggal R, Hao W. 2008. Novel chimeric genotype 1b/2a hepatitis C virus suitable for high-throughput screening. Antimicrob Agents Chemother 52:666–674.

21. Jones CT, Murray CL, Eastman DK, Tassello J, Rice CM. 2007. Hepatitis C virus p7 and NS2 proteins are essential for production of infectious virus. J Virol 81:8374–8383.

22. Horwitz JA, Dorner M, Friling T, Donovan BM, Vogt A, Loureiro J, Oh T, Rice CM, Ploss A. 2013. Expression of heterologous proteins flanked by NS3-4A cleavage sites within the hepatitis C virus polyprotein. Virology 439:23–33.

23. Counihan NA, Rawlinson SM, Lindenbach BD. 2011. Trafficking of hepatitis C virus core protein during virus particle assembly. PLoS Pathog 7:e1002302.

24. Tamura T, Fukuhara T, Uchida T, Ono C, Mori H, Sato A, Fauzyah Y, Okamoto T, Kurosu T, Setoh YX, Imamura M, Tautz N, Sakoda Y, Khromykh AA, Chayama K, Matsuura Y. 2018. Characterization of Recombinant Flaviviridae Viruses Possessing a Small Reporter Tag. J Virol 92:e01582–17.

25. Moradpour D, Evans MJ, Gosert R, Yuan Z, Blum HE, Goff SP, Lindenbach BD, Rice CM. 2004. Insertion of green fluorescent protein into nonstructural protein 5A allows direct visualization of functional hepatitis C virus replication complexes. J Virol 78:7400–7409.

26. Gottwein JM, Jensen TB, Mathiesen CK, Meuleman P, Serre SBN, Lademann JB, Ghanem L, Scheel TKH, Leroux-Roels G, Bukh J. 2011. Development and application of hepatitis C reporter viruses with genotype 1 to 7 core-nonstructural protein 2 (NS2) expressing fluorescent proteins or luciferase in modified JFH1 NS5A. J Virol 85:8913–8928.

27. Watashi K, Hijikata M, Hosaka M, Yamaji M, Shimotohno K. 2003. Cyclosporin A suppresses replication of hepatitis C virus genome in cultured hepatocytes. Hepatology 38:1282–1288.

28. Paeshuyse J, Kaul A, De Clercq E, Rosenwirth B, Dumont J-M, Scalfaro P, Bartenschlager R, Neyts J. 2006. The non-immunosuppressive cyclosporin DEBIO-025 is a potent inhibitor of hepatitis C virus replication in vitro. Hepatology 43:761–770.

29. Lawitz E, Mangia A, Wyles D, Rodriguez-Torres M, Hassanein T, Gordon SC, Schultz M, Davis MN, Kayali Z, Reddy KR, Jacobson IM, Kowdley KV, Nyberg L, Subramanian GM, Hyland RH, Arterburn S, Jiang D, McNally J, Brainard D, Symonds WT, McHutchison JG, Sheikh AM, Younossi Z, Gane EJ. 2013. Sofosbuvir for previously untreated chronic hepatitis C infection. N Engl J Med 368:1878–1887.

30. Stuyver LJ, Whitaker T, McBrayer TR, Hernandez-Santiago BI, Lostia S, Tharnish PM, Ramesh M, Chu CK, Jordan R, Shi J, Rachakonda S, Watanabe KA, Otto MJ, Schinazi RF. 2003. Ribonucleoside analogue that blocks replication of bovine viral diarrhea and hepatitis C viruses in culture. Antimicrob Agents Chemother 47:244–254.

31. Arumugaswami V, Remenyi R, Kanagavel V, Sue EY, Ngoc Ho T, Liu C, Fontanes V, Dasgupta A, Sun R. 2008. High-resolution functional profiling of hepatitis C virus genome. PLoS Pathog 4:e1000182.

32. Wang C, Le SY, Ali N, Siddiqui A. 1995. An RNA pseudoknot is an essential structural element of the internal ribosome entry site located within the hepatitis C virus 5’ noncoding region. RNA 1:526–537.

33. Lohmann V, Körner F, Koch J, Herian U, Theilmann L, Bartenschlager R. 1999. Replication of subgenomic hepatitis C virus RNAs in a hepatoma cell line. Science 285:110–113.

34. Jirasko V, Montserret R, Appel N, Janvier A, Eustachi L, Brohm C, Steinmann E, Pietschmann T, Penin F, Bartenschlager R. 2008. Structural and functional characterization of nonstructural protein 2 for its role in hepatitis C virus assembly. J Biol Chem 283:28546–28562.

35. de la Fuente C, Goodman Z, Rice CM. 2013. Genetic and functional characterization of the N-terminal region of the hepatitis C virus NS2 protein. J Virol 87:4130–4145.

36. Scheel TKH, Prentoe J, Carlsen THR, Mikkelsen LS, Gottwein JM, Bukh J. 2012. Analysis of functional differences between hepatitis C virus NS5A of genotypes 1-7 in infectious cell culture systems. PLoS Pathog 8:e1002696.

37. Gaspar N, Zambito G, Dautzenberg IJC, Cramer SJ, Hoeben RC, Lowik C, Walker JR, Kirkland TA, Smith TP, van Weerden WM, de Vrij J, Mezzanotte L. 2020. NanoBiT system and hydrofurimazine for optimized detection of viral infection in mice-A novel in vivo imaging platform. Int J Mol Sci 21:5863.

38. Kim JH, Bryant H, Fiedler E, Cao T, Rayner JO. 2022. Real-time tracking of bioluminescent influenza A virus infection in mice. Sci Rep 12:3152.

39. Gallardo-Flores CE, Colpitts CC. 2021. Cyclophilins and their roles in hepatitis C virus and Flavivirus infections: Perspectives for novel antiviral approaches. Pathogens 10:902.

40. Coelmont L, Hanoulle X, Chatterji U, Berger C, Snoeck J, Bobardt M, Lim P, Vliegen I, Paeshuyse J, Vuagniaux G, Vandamme A-M, Bartenschlager R, Gallay P, Lippens G, Neyts J. 2010. DEB025 (Alisporivir) inhibits hepatitis C virus replication by preventing a cyclophilin A induced cis-trans isomerisation in domain II of NS5A. PLoS One 5:e13687.

41. Page K, Melia MT, Veenhuis RT, Winter M, Rousseau KE, Massaccesi G, Osburn WO, Forman M, Thomas E, Thornton K, Wagner K, Vassilev V, Lin L, Lum PJ, Giudice LC, Stein E, Asher A, Chang S, Gorman R, Ghany MG, Liang TJ, Wierzbicki MR, Scarselli E, Nicosia A, Folgori A, Capone S, Cox AL. 2021. Randomized Trial of a Vaccine Regimen to Prevent Chronic HCV Infection. N Engl J Med 384:541–549.

42. Lindow M, Kauppinen S. 2012. Discovering the first microRNA-targeted drug. J Cell Biol 199:407–412.

43. Rozewicki J, Li S, Amada KM, Standley DM, Katoh K. 2019. MAFFT-DASH: integrated protein sequence and structural alignment. Nucleic Acids Res 47:W5–W10.

